# Biochemical ‘Cambrian’ explosion-implosions: the generation and pruning of genetic codes

**DOI:** 10.1101/002436

**Authors:** Rodrick Wallace

## Abstract

Tlusty’s topological analysis of the genetic code suggests ecosystem changes in available metabolic free energy that predated the aerobic transition enabled a punctuated sequence of increasingly complex genetic codes and protein translators. These coevolved via a ‘Cambrian explosion’ until, very early on, the ancestor of the present narrow spectrum of protein machineries became evolutionarily locked in at a modest level of fitness reflecting a modest embedding metabolic free energy ecology. Similar biochemical ‘Cambrian singularities’ must have occurred at different scales and levels of organizaron on Earth, with competition or chance-selected outcomes frozen at a far earlier period than the physical bauplan Cambrian explosion. Other examples might include explosive variations in mechanisms of photosynthesis and subsequent oxygen metabolisms. Intermediate between Cambrian bauplan and genetic code, variants of both remain today, even after evolutionary pruning, often protected in specialized ecological niches. This suggests that, under less energetic astrobiological ecologies, a spectrum of less complicated reproductive codes may also survive in specialized niches.

## 1 Introduction

Wallace [1] argues that ‘Cambrian explosions’ are standard features of blind evolutionary process, representing outliers in the ongoing routine of evolutionary punctuated equilibrium. Most such explosions, however, will be severely pruned by selection and chance extinction. That work suggested, in passing, that the evolution of the genetic code, involving the transmission of information between codon machinery and amino acid machinery, is likely to have undergone just such an ‘explosion’ as significant levels of chemical free energy became available to metabolic process. Here, we make that argument explicit.

See figure 1, taken from [2], for a schematic of the present highly evolved system relating code to protein component. In modern protein synthesis, the anticodon at one end of a tRNA molecule binds to its complementary codon in mRNA derived directly from the genome. Sequence-to-sequence translation is not highly parallel, in this model, and the process can be characterized in terms of the Shannon uncertainty in the transmission of information between codon machinery and amino acid machinery.

**Figure 1:**
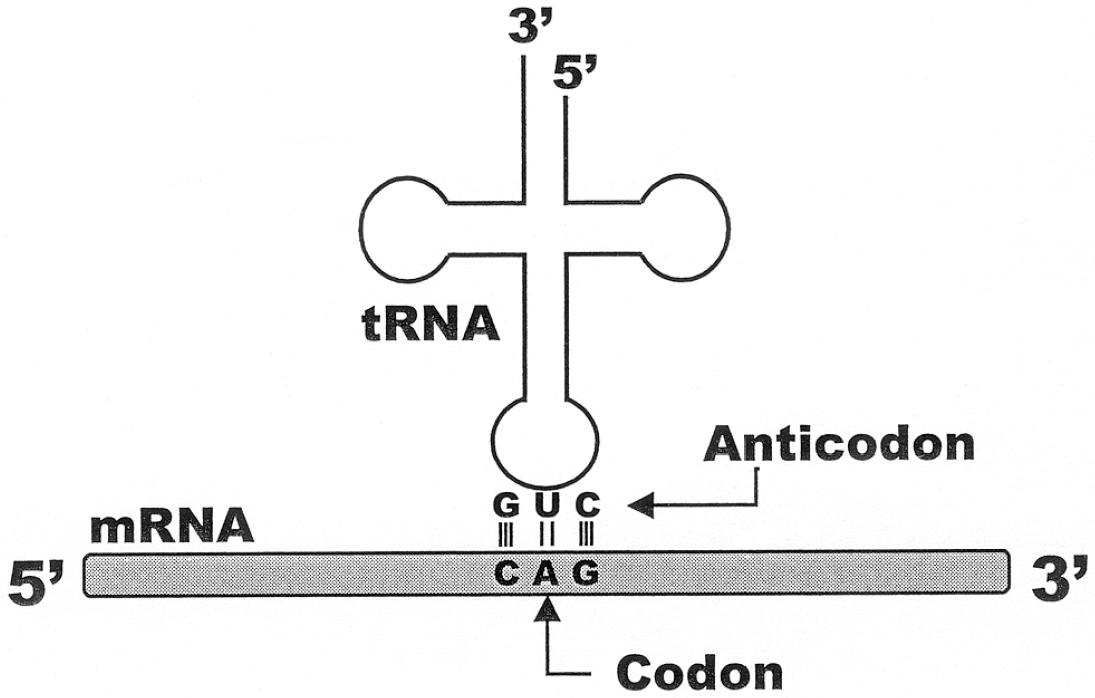
Adapted from fig. 1.8 of [2]. Modern protein synthesis; the anticodon at one end of a tRNA molecule binds to its complementary codon in mRNA derived directly from the genome. Sequence-to-sequence translation is not highly parallel, in this model, and the process can be characterized in terms of the Shannon uncertainty in the transmission of information between codon machinery and amino acid machinery.

To paraphrase Tlusty [3], the genetic code emerges as a transition in a noisy information channel, using the Rate Distortion Theorem: the optimal code is described by the minimum of a ‘free energy’-like functional, which leads naturally to the possibility of describing the code’s emergence as a transition akin to a phase transition in statistical physics. The basis for this is the observation that a supercritical phase transition is known to take place in noisy information channels (e.g., [4]). The noisy channel is controlled by a temperature-like parameter that determines the balance between the information rate and the distortion ‘in the same way that physical temperature controls the balance between energy and entropy’ in a physical system. Following Tlusty’s equation (2), the ‘free energy’ functional has the form *D* − *TS* where *D* is the average ‘error load’, equivalent to average distortion in a rate distortion problem, *S* is the ‘entropy due to random drift’, and *T* measures the strength of random drift relative to the selection force that pushes towards fitness maximization. This is essentially a Morse function, in the sense of the Mathematical Appendix. According to Tlusty’s analysis, at high *T* the channel is totally random and it conveys zero information. At a certain critical temperature *T_c_* the information rate starts to increase continuously. In subsequent work, Tlusty [5] expands the analysis to include the interplay of accuracy, diversity, and the cost of the coding machinery.

More specifically, as Tlusty [3] puts it,

> To discuss the topology of errors we portray the codon space as a graph whose verticies are the codons… Two codons… are linked by an edge if they are likely to be confused by misreading… We assume that two codons are most likely to be confused if all their letters except for one agree and therefore draw an edge between them. The resulting graph is natural for considering the impact of translation errors on mutations because such errors almost always involve a single letter difference, that is, a movement along an edge of the graph to a neighboring vertex.
>
> The topology of a graph is characterized by its genus *γ*, the minimal number of holes required for a surface to embed the graph such that no two edges cross. The more connected that a graph is the more holes are required for its minimal embedding… [T]he highly interconnected 64-codon graph is embedded in a holey, *γ* = 41 surface. The genus is somewhat reduced to *γ* = 25 if we consider only 48 effective codons.

Tlusty further concludes that the topology of the code sets an upper limit to the number of low modes – critical points – of his free energy-analog functional, and this is also the number of amino acids. The low modes define a partition of the codon surface into domains, and in each domain a single amino acid is encoded. The partition optimizes the average distortion by minimizing the boundaries between the domains as well as the dissimilarity between neighboring amino acids.

Tlusty states:

> The maximum [of the functional] determines a single contiguous domain where a certain amino acid is encoded… Thus every mode corresponds to an amino acid and the number of modes is the number of amino acids. This compact organization is advantageous because misreading of one codon as another codon within the same domain has no deleterious impact. For example, if the code has two amino acids, it is evident that the error-load of an arrangement where there are two large contiguous regions, each coding for a different amino acid, is much smaller than a ‘checkerboard’ arrangement of the amino acids.

This, Tlusty [6] points out, is analogous to the well-known topological coloring problem: “in the coding problem one desires maximal similarity in the colors of neighboring ‘countries’, while in the coloring problem one must color neighboring countries by different colors”. After some development [5], the number of possible amino acids in this scheme is determined by Heawood’s formula [7]:

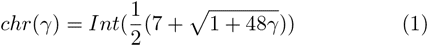

where *chr*(*γ*) is the number of color domains of a surface with genus *γ*, and *Int*(*x*) is the integer value of *x*.

However, from Morse Theory [8]:

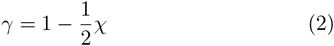

where *χ* is the Euler characteristic of the underlying topological manifold. For a manifold having a Morse function *f*, *χ* can be expressed as the alternating sum of the function’s Morse numbers: The Morse numbers *µ_i_*(*i* = 0,1, …, *m*) of *f* on the manifold are the number of critical points (*df* (*x*_*c*_) = 0) of index *i*, the number of negative eigenvalues of the matrix 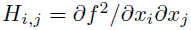. Then 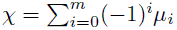.

This holds true for any Morse function on the manifold *M*. See the Mathematical Appendix for a summary of material on Morse Theory.

We reproduce part of Table 1 of [3], showing the topological limit to the number of amino acids for different codes:

| Code | # Codons | Max. # AA’s |
| --- | --- | --- |
| 4-base singlets | 4 | 4 |
| 3-base doublets | 9 | 7 |
| 4-base doublets | 16 | 11 |
| 16 codons | 32 | 16 |
| 48 codons | 48 | 20 |
| 4-base triplets | 64 | 25 |

This is the fundamental topological decomposition, to which Morse-theoretic ‘free energy’ functionals are to be fit. Note, however, that, while the scheme limits underlying code bauplan, for the current coding a simple combinatorial argument shows there are 10^84^ possible alternative code tables if each of the 20 amino acids and the stop signal are assigned at least one codon. Smaller, but still astronomical, numbers can be associated with the less complicated codes, permitting a later statistical mechanics-like model driven by available metabolic free energy.

Tlusty [3] concludes:

> [This] suggests a pathway for the evolution of the present-day code from simpler codes, driven by the increasing accuracy of improving translation machinery. Early translation machinery corresponds to smaller graphs since indiscernible codons are described by the same vertex. As the accuracy improves these codons become discernible and the corresponding vertex splits. This gives rise to a larger graph that can accommodate more amino acids… [P]resent-day translation machinery with a four-letter code and 48–64 codons (no discrimination between U and C in the third position) gave rise to 20–25 amino acids. One may think of future improvement that will remove the ambiguity in the third position (64 discernible codons). This is predicted to enable stable expansion of the code up to 25 amino acids.

The underlying model is that of phase transitions in physical systems. Following Landau’s group symmetry shifting arguments [9, 10], higher temperatures enable higher system symmetries, and, as temperature changes, punctuated shifts to different symmetry states that occur in characteristic manners. Extension of this argument in terms of information transmission between codons and proteins, in the context of metabolic energy measures, seems direct, particularly involving the groupoids constructed by the disjoint union of the homology groups representing the different coding topologies that Tlusty identifies. A more complete mathematical treatment of some of these and related matters can be found in [11, 12].

It is worth noting that Tlusty’s method can be applied as well to large-scale globular protein folding [13]. Equation (1), most simply, produces the table

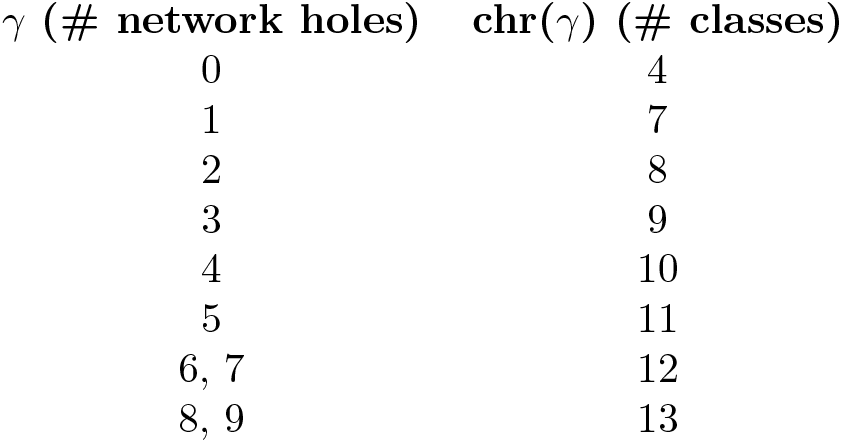

In Tlusty’s scheme, the second column represents the maximal possible number of product classes that can be reliably produced by error-prone codes having γ holes in the underlying coding error network.

Normal irregular protein symmetries were first classified by Levitt and Chothia [14], following a visual study of polypeptide chain topologies in a limited dataset of globular proteins. Four major classes emerged; all α-helices; all *β*-sheets; *α*/*β*; and *α* + *β*, with the latter two having the obvious meaning.

While this scheme strongly dominates observed irregular protein forms, Chou and Maggiora [15], using a much larger data set, recognize three more ‘minor’ symmetry equivalence classes; *µ* (multi-domain); *σ* (small protein); and *ρ* (peptide), and a possible three more subminor groupings.

We infer that the normal globular ‘protein folding code error network’ is, essentially, a large connected ‘sphere’ – producing the four dominant structural modes – having one minor, and possibly as many as three more subminor attachment handles, in the Morse Theory sense [8], a matter opening up other analytic approaches.

From Tlusty’s perspective, then, four-fold protein folding classification produces the simplest possible large-scale ‘protein folding code’, a sphere limited by the four-color problem, and the simplest cognitive cellular regulatory system would thus be constrained to pass/fail on four basic flavors, as it were, of folded proteins.

Here we will reconsider the evolutionary trajectories of genetic codes in the context of intensive measures of available metabolic free energy, taking the perspective that the availability of metabolic free energy is central to the evolution of complex phenomena of biological communication. That is, the ‘temperature’ analog is the chemical free energy available to early anaerobic metabolisms, in the sense of Canfield et al. [16]. For example, the proposed hydrogen/sulfur reaction

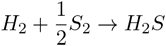

produces something like *M* = 21 KJ/mol, while the aerobic reaction

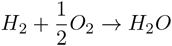

produces about *M* = 241 KJ/mol, more than an order of magnitude greater. The genetic code, however, was locked in by evolutionary path dependence well before oxygen became widely available for metabolic process.

Figure 2, taken from [16] shows a range of possible electron donors and acceptors available to early anaerobic metabolisms on earth.

**Figure 2:**
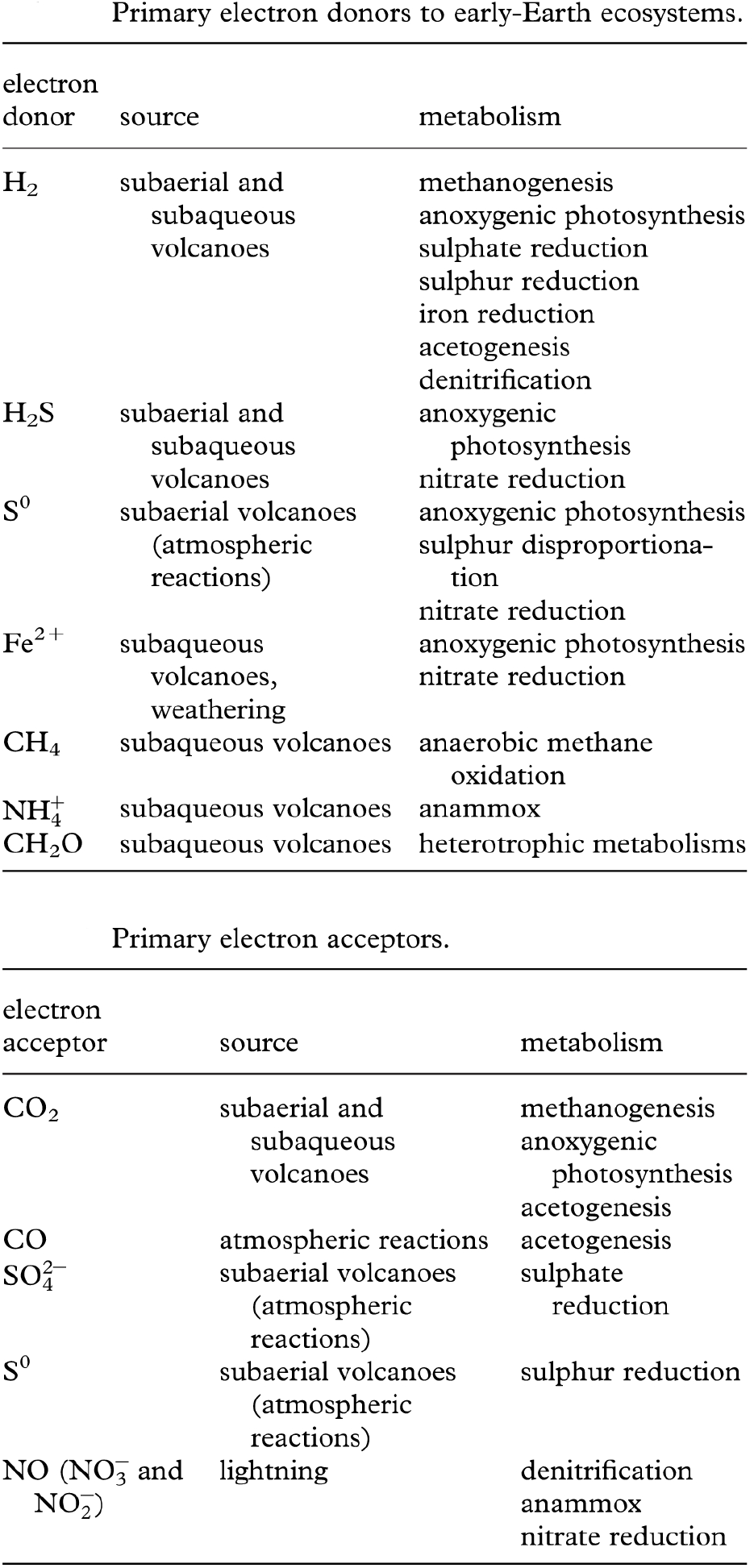
From Canfield et al., [16], Possible electron donors and acceptors for early-Earth ecosystems.

We begin with a restatement of a few central ideas from [1].

## 2 Information theory

A genetic code, translating codons into proteins, implies the existence of an information source using that code, and the behavior of such sources is constrained by the asymptotic limit theorems of information theory. Thus, the interaction between biological subsystems associated with a code can be formally restated in communication theory terms. Wallace and Wallace [17, 18] use an elaborate cognitive paradigm for gene expression to infer such information sources, i.e., cognition implies ‘language’, in a large sense, but the focus here on codes condenses the argument because a code directly implies existence of an information source using it.

Here we think of the machinery listing a sequence of codons as communicating with machinery that produces amino acids, allowing definition of an information source embedded in an environment whose regularities themselves imply the existence of an information source.

Following [1], assume there are *n* possible genetic code ‘species’ interacting with an embedding environment represented by an information source *Z*. The processes associated with each code species *i* are represented as information sources *X_i_*. These information sources undergo a ‘coevolutionary’ interaction in the sense of [19], producing a joint information source uncertainty [20] for the full system as

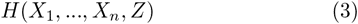

Feynman’s [21] insight that information is a form of free energy allows definition of an entropy-analog as

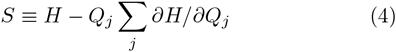

The *Q_i_* are taken as driving parameters that may include, but are not limited to, the Shannon uncertainties of the underlying information sources.

Again following Wallace (2014), we can characterize the dynamics of the system in terms of onsager-like nonequilibrium thermodynamics in the gradients of *S* as the set of stochastic differential equations [22],

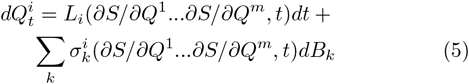

where the *B_k_* represent noise terms having particular forms of quadratic variation. See [23] or other standard references on stochastic differential equations for details.

This can be more simply written as

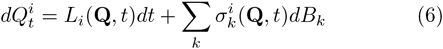

where **Q** ≡ (*Q*^1^,…,*Q*^*m*^)

Following the arguments of Champagnat et al., this is a coevolutionary structure, where fundamental dynamics are determined by component interactions:

1. Setting the expectation of equations (5) equal to zero and solving for stationary points gives attractor states since the noise terms preclude unstable equilibria. These are analogous to the evolutionarily stable states of evolutionary game theory.
2. This system may, however, converge to limit cycle or pseudorandom ‘strange attractor’ behaviors similar to thrashing in which the system seems to chase its tail endlessly within a limited venue – the ‘Red Queen’.
3. What is ‘converged’ to in any case is not a simple state or limit cycle of states. Rather it is an equivalence class, or set of them, of highly dynamic information sources coupled by through crosstalk and other mutual interactions. Thus ‘stability’ in this structure represents particular patterns of ongoing dynamics rather than some identifiable static configuration, that is, at best, a nonequilibrium steady state.
4. Applying Ito’s chain rule for stochastic differential equations to the 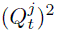 and taking expectations allows calculation of variances. These may depend very powerfully on a system’s defining structural constants, leading to significant instabilities [24].

## 3 Large deviations: iterating the model

As Champagnat et al. note, shifts between the quasi-equilibria of a coevolutionary system can be addressed by the large deviations formalism. The dynamics of drift away from trajectories predicted by the canonical equation can be investigated by considering the asymptotic of the probability of ‘rare events’ for the sample paths of the diffusion.

‘Rare events’ are the diffusion paths drifting far away from the direct solutions of the canonical equation. The probability of such rare events is governed by a large deviation principle, driven by a ‘rate function’ 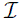 that can be expressed in terms of the parameters of the diffusion.

This result can be used to study long-time behavior of the diffusion process when there are multiple attractive singularities. Under proper conditions, the most likely path followed by the diffusion when exiting a basin of attraction is the one minimizing the rate function 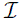 over all the appropriate trajectories.

An essential fact of large deviations theory, however, is that the rate function 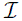 almost always has the canonical form

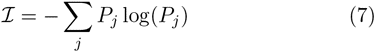

for some probability distribution, i.e., the uncertainty of an information source [25].

The argument directly complements equation (5), now seen as subject to large deviations that can themselves be described as the output of an information source *L_D_* defining 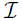, driving or defining *Q^j^*-parameters that can trigger punctuated shifts between quasi-stable nonequilibrium steady states.

Not all large deviations are possible: only those consistent with the high probability paths defined by the information source *L_D_* will take place.

Recall from the Shannon-McMillan Theorem [26] that the output streams of an information source can be divided into two sets, one very large that represents nonsense statements of vanishingly small probability, and one very small of high probability representing those statements consistent with the inherent ‘grammar’ and ‘syntax’ of the information source. For example, whatever higher-order multicellular evolution takes place, some equivalent of backbone and blood remains.

Thus we could now rewrite equation (3) as

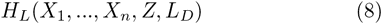

where we have explicitly incorporated the ‘large deviations’ information source *L_D_* that defines high probability evolutionary excursions for this system.

Again carrying out the argument leading to equation (5), we arrive at another set of quasi-stable modes, but possibly very much changed in number; either branched outward in time by a wave of speciation or quasi-speciation, or decreased through a wave of extinction. Iterating the models backwards in time constitutes a cladistic or coalescent analysis.

## 4 Evolution of genetic codes under relaxed path-dependence

Following the arguments of [1], in general, for current organisms, the number of quasi-equilibria available to the system defined by equation (5), or to its generalization via equation (7), will be quite small — indeed, at most a handful — a consequence of code lock-in by path dependent evolutionary process. The same cannot be said, however, for earlier species or quasi-species, to which can be applied more general methods that may represent key processes acting three or four billion years in the past.

Under such a relaxation assumption, the speciation/extinction large deviations information source *L_D_* is far less constrained, and there will be very many possible quasistable nonequilibrium steady states available for transition, analogous to an ensemble in statistical mechanics. Again, for the current genetic code, involving 20 possible amino acids, following the arguments of Table 1, there are some 10^84^ possible alternative codings.

The metabolic free energy index in KJ/mol, which we write as *M*, can, from the arguments of the Introduction, then be interpreted as a kind of temperature measure so that higher values permit higher equivalence class groupoid symmetries. This leads to a relatively simple statistical mechanics analog built on the *H_L_* of equation (7).

Define a pseudoprobability for quasi-stable mode *j* as

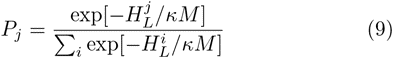

where κ is a scaling constant and *M* is a reaction energy intensity, typically measured as KJ/mol.

Next, define a Morse Function *F* as

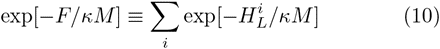

Apply Pettini’s topological hypothesis to *F*. Then *M* is seen as a very general temperature-like intensity measure whose changes drive punctuated topological alterations in the underlying ecological and coevolutionary structures associated with the Morse Function *F*.

Such topological changes, following Pettini’s arguments, can be far more general than indexed by the simple Landau-type critical point phase transition in an order parameter.

Thus, the results of [27], regarding the complexity of the genetic code, can be directly reframed in terms of available metabolic free energy intensity leading to equations (9) and (10). Then *M* is a measure of metabolic free energy intensity, and the 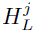 represent the Shannon uncertainties in the transmission of information between codon machinery and amino acid machinery.

Increasing *M* then leads to the possibility of more complex genetic codes, i.e., those having higher measures of symmetry, in the Landau sense, as calculated by Tlusty’s methods, until competition, selection, and chance extinction leading to evolutionary lock-in took place at a relatively low level of coding efficiency. Canfield et al. [16] speculate that the most active early ecosystems were probably driven by the cycling of *H*_2_ and *Fe*^2+^, providing relatively low free energy intensities for metabolic process.

## 5 Discussion and conclusions

Marshall [28] characterizes the ‘Cambrian explosion’ in animal physical bauplan that took place 500 myr ago as follows:

> With the advent of ecological interactions between macroscopic adults… especially… predation…, the number of needs each organism had to meet must have increased markedly: Now there were myriad predators to contend with, and a myriad number of ways to avoid them, which in turn led to more specialized ways of predation as different species developed different avoidance strategies, etc… The combinatoric richness already present in the Ediacaran genome was extracted through the richness of biotic interaction as the Cambrian ‘explosion’ unfolded…

Here we argue that, in analogous fashion, the availability of myriad biochemical electron donor/acceptor cycles according to the schema of [16] in figure 2 created a rich chemical ecology. The explosive combinatoric richness present in the possible variety of genetic codes was, however, in this case extracted through the richness of biotic interaction downward in scale by the efficiency, effectiveness, or chance survival, of a single code that emerged at a modest level of available chemical free energy to persist as the present dominant genetic code.

Wallace [29] makes a similar argument for the evolution of homochirality, formally invoking the standard groupoid approach to stereochemistry in a thermodynamic context that likewise generalizes Landau’s spontaneous symmetry breaking arguments. On Earth, limited metabolic free energy density may have served as a low temperature-analog to ‘freeze’ the system in the lowest energy state, i.e., the set of simplest homochiral transitive groupoids representing reproductive chemistries. These engaged in Darwinian competition until a single configuration survived. Subsequent path-dependent evolutionary process locked-in this initial condition. Astrobiological outcomes, in the presence of higher initial metabolic free energy densities, could well be considerably richer, for example, of mixed chirality. One result would be a complicated distribution of biological chirality across a statistically large sample of extraterrestrial stereochemistry, in marked contrast with recent published analyses predicting a racemic average.

The essential insight is that ‘Cambrian singularities’ at different scales and levels of organization are inherently path dependent, with the evolutionarily or chance-selected outcomes of basic biochemical explosions likely to be locked in at a far earlier period than physical bauplan explosions. Possible examples include explosive evolutionary variations in photosynthesis and mechanisms of tissue oxidation. Different forms of both remain today, even after evolutionary pruning, particularly variants in the ‘bauplan’ of oxidizer metabolism (e.g., [30, 31]).

This suggests that, under less energetic astrobiological ecologies, a spectrum of less complex reproductive codes may survive, likely across a set of characteristic ecosystem niches.

## 6. Acknowledgments

The author thanks Dr. D.N. Wallace for useful discussions.

## 7 Mathematical Appendix

Morse theory examines relations between analytic behavior of a function – the location and character of its critical points – and the underlying topology of the manifold on which the function is defined. Here we follow closely [10].

The essential idea of Morse theory is to examine an *n*-dimensional manifold *M* as decomposed into level sets of some function *f* : *M* → **R** where **R** is the set of real numbers. The *a*-level set of *f* is defined as

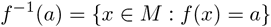

the set of all points in *M* with *f* (*x*) = *a*. If *M* is compact, then the whole manifold can be decomposed into such slices in a canonical fashion between two limits, defined by the minimum and maximum of *f* on *M*. Let the part of *M* below *a* be defined as

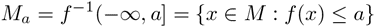

These sets describe the whole manifold as *a* varies between the minimum and maximum of *f*.

Morse functions are defined as a particular set of smooth functions *f* : *M* → **R** as follows. Suppose a function *f* has a critical point *x*_c_, so that the derivative *df* (*x*_*c*_) = 0, with critical value *f* (*x*_*c*_). Then *f* is a Morse function if its critical points are nondegenerate in the sense that the Hessian matrix of second derivatives at *x*_*c*_, whose elements, in terms of local coordinates are given by *H_i,j_* = ∂^2^*f*/∂*x^i^*∂*x^j^*, has rank *n*, which means that it has only nonzero eigenvalues, so that there are no lines or surfaces of critical points and, ultimately, critical points are isolated.

The index of the critical point is the number of negative eigenvalues of *H* at *x*_*c*_.

A level set *f*^−1^(a) of f is called a critical level if *a* is a critical value of *f*, that is, if there is at least one critical point *x*_*c*_ ∈ *f*^−1^(*a*).

The essential results of Morse theory are:

1. If an interval [*a*, *b*] contains no critical values of *f*, then the topology of *f*^−1^[*a*, *v*] does not change for any 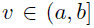. Importantly, the result is valid even if *f* is not a Morse function, but only a smooth function.
2. If the interval [*a, b*] contains critical values, the topology of *f*^−1^ [*a*, *v*] changes in a manner determined by the properties of the matrix *H* at the critical points.
3. If f: *M* → **R** is a Morse function, the set of all the critical points of *f* is a discrete subset of *M*, i.e., critical points are isolated. This is Sard’s Theorem.
4. If f: *M* → **R** is a Morse function, with *M* compact, then on a finite interval [*a*, *b*] ⊂ **R**, there is only a finite number of critical points *p* of *f* such that *f* (p) ∈ [*a*, *b*]. The set of critical values of *f* is a discrete set of **R**.
5. For any differentiable manifold *M*, the set of Morse functions on *M* is an open dense set in the set of real functions of *M* of differentiability class *r* for 0 ≤ *r* ≤ ∞.
6. Some topological invariants of *M*, that is, quantities that are the same for all the manifolds that have the same topology as *M*, can be estimated and sometimes computed exactly once all the critical points of f are known: Let the Morse numbers *µ*_*i*_ (*i* = 0,…, *m*) of a function f on *M* be the number of critical points of *f* of index *i*, (the number of negative eigenvalues of *H*). The Euler characteristic of the complicated manifold *M* can be expressed as the alternating sum of the Morse numbers of any Morse function on *M*,

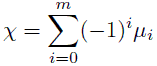

The Euler characteristic reduces, in the case of a simple polyhedron, to

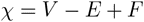

where *V, E*, and *F* are the numbers of vertices, edges, and faces in the polyhedron.
7. Another important theorem states that, if the interval [*a, b*] contains a critical value of *f* with a single critical point *x_c_*, then the topology of the set *M*_*b*_ defined above differs from that of *M_a_* in a way which is determined by the index, *i*, of the critical point. Then *M*_*b*_ is homeomorphic to the manifold obtained from attaching to *M*_*a*_ an *i*-handle, i.e. the direct product of an *i*-disk and an *(m — i*)-disk.

Again, [8, 10] contain both mathematical details and further references.

